# A ballistic pollen dispersal strategy hidden in stylar oscillation

**DOI:** 10.1101/2020.01.27.920710

**Authors:** Shuto Ito, Hamed Rajabi, Stanislav N Gorb

## Abstract

Asteraceae, the most successful flowering plant family, is adapted to the vast range of ecological niches. Their adaptability is partially based on their strong ability of reproduction. The initial, yet challenging, step for plant reproduction is to transport pollen to flower-visiting pollinators. Using quantitative experiments and numerical simulations, here we show that the common floral feature of Asteraceae, a pollen-bearing style, serves as a ballistic lever for catapulting pollen grains to pollinators. This is likely to be a pollination strategy to propel pollen to blind spots of pollinators’ bodies, which are beyond the physical reach of the styles. Our results suggest that the specific morphology and length of the floret, as well as the pollen adhesion, avoid pollen waste by catapulting pollen within a certain range equal to the size of a flowerhead. The insights into the functional floral oscillation may shed light on the superficially unremarkable, but ubiquitous functional floral design of Asteraceae.

## 1. Introduction

Pollination, the journey of pollen grains from an anther to a receptive stigma of another plant, is a key to successful plant reproduction. Vast majority of flowering plants rely on pollinators for pollen transport^1,2^. Hence, the design of floral systems that enables the efficient pollen transport via the interaction with pollinators can determine the reproductive success of species.

The most successful flowering plant family Asteraceae, with roughly 10% of the total number of flowering plants, inhabit every continent except Antarctica^3^. Asteraceae are distinguishable by their clusters of numerous individual flowers, called florets (Fig. 1). Pollination in Asteraceae is initiated by releasing pollen grains into anther tubes. As styles elongate within the anther tubes, they push or brush the pollen out of the anther tubes, exposing the pollen grains at the stylar surface^4,5^.

**Fig. 1.**
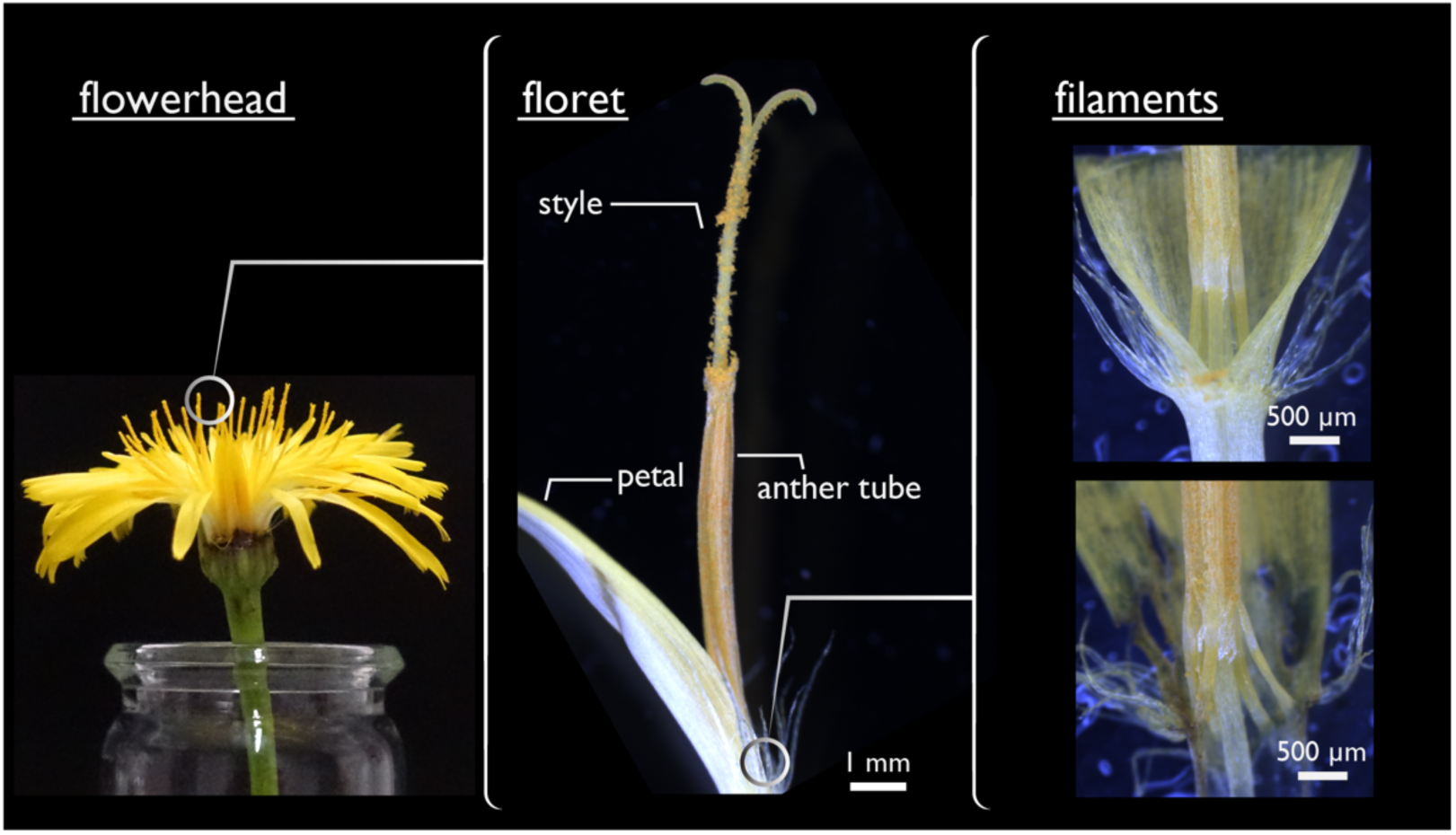
Floral structures of *Hypochaeris radicata* at three different scales: a whole flowerhead (left) composed of many florets (middle), of which basal segment, so called filaments, are magnified (right).

According to their preferences for pollinators, Asteraceae are known as generalists. This means that exposed pollen grains on their styles are pollinated by various insect groups including Coleoptera, Diptera, and Hymenoptera^6^. Hence, it is likely that Asteraceae have secured pollination by using a variety of strategies, which fit to the wide range of body shapes and sizes of their diverse pollinators.

An intuitive pollination strategy is achieved by direct physical contact between exposed pollen grains and visiting pollinators (Fig. 2a). This strategy, however, has a few disadvantages. First, some pollinators, such as bees, visit flowers to feed on pollen grains. Therefore, pollen that are actively collected by such pollinators are usually wasted. Second, some pollinators utilize their elongated mouth parts to steal nectars without frequent contacts with exposed pollen grains (Fig. 2f-h). Therefore, in such cases, no or only a small amount of pollen is transferred, which might be insufficient for successful pollination. How do Asteraceae compensate for the aforementioned shortcomings?

**Fig. 2.**
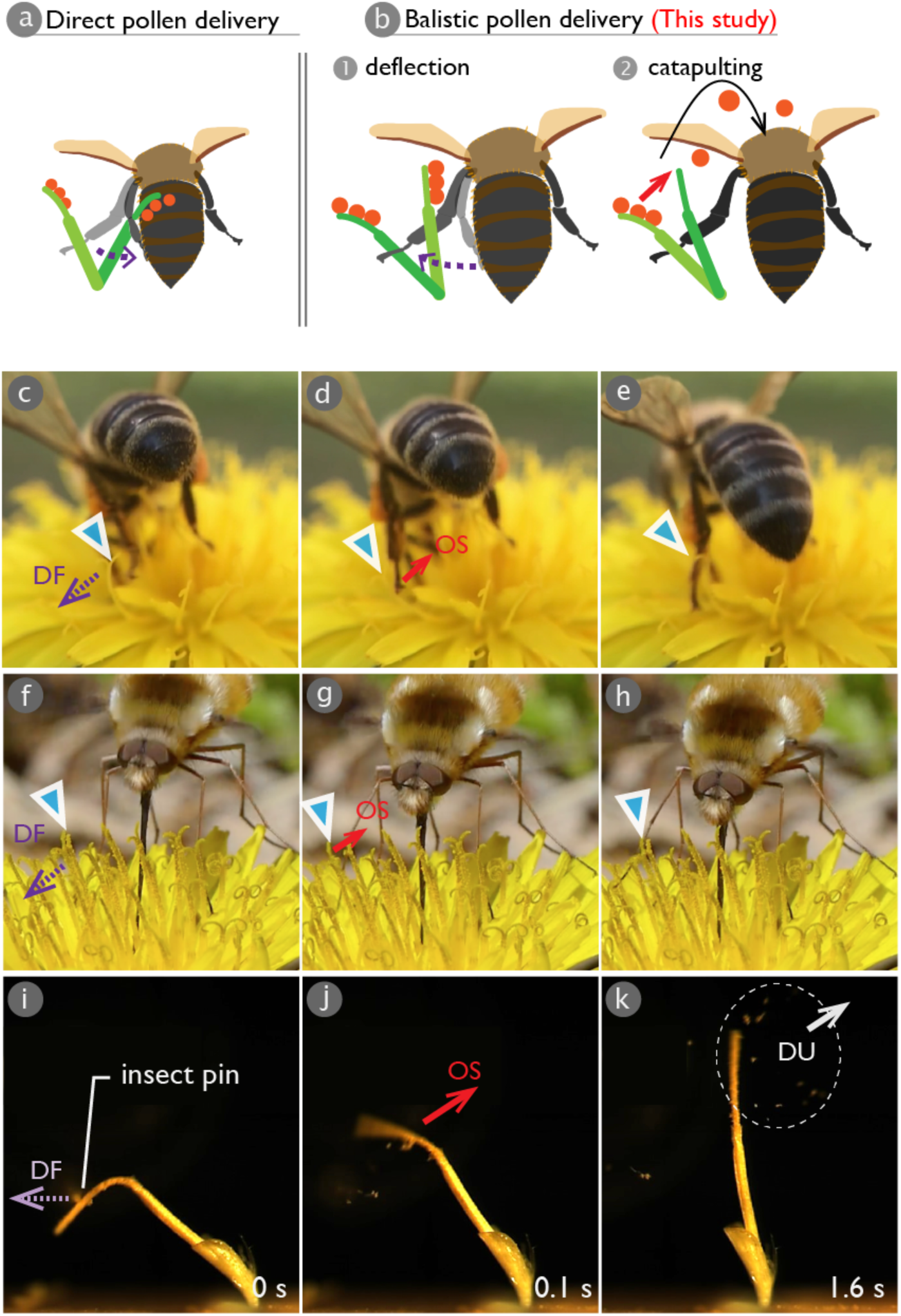
(a,b) Conceptualization of the current study. A stylar deflection toward a pollinator causes a direct pollen delivery to the pollinator (a), whereas a deflection to the opposite direction enables a ballistic pollen delivery to the pollinator (b). (c-h) Interaction between insects and styles. A style, which position is shown with blue triangle, was deflected by a limb of a bee *Apis mellifera* (c-e) or that of a fly from the family Bombyliidae (f-h). Upon the release, the style snaps back towards the insect. (d-f) Snapshots from a high-speed video of an oscillating style showing pollen dispersal after a larger deflection. DF: deflection, OS: oscillation, DU: pollen dispersal units.

Electrostatic attraction is known to facilitate pollen transport to pollinators^7–9^. However, its efficiency strongly drops under high humidity levels^10^. Buzz pollination, another potential strategy, in which pollen are released from flowers using vibrations caused by bees and bumblebees, is exclusive to only very specialized plant species^11,12^, and therefore, it is not the case for the “generalist” Asteraceae.

Considering that the habitat requirements of Asteraceae are met in various locations, it is likely that their alternative pollination strategy (1) has versatile working principles to enable successful adaption in various climates and ecologies, (2) is triggered without frequent physical contacts between pollen and pollinators, and (3) enables the attachment of pollen to the “blind spots” of pollinators^13,14^, where they cannot reach during active grooming. Here, based on our quantitative experiments and numerical simulations, we propose a previously unexplored pollen dispersal strategy in Asteraceae, which can satisfy all the requirements mentioned above.

## 2. Material and methods

### 2.1. Plant Species

*Hypochaeris raticata* (Asteraceae) was previously adopted as a model species to investigate pollen adhesion^15,16^. In this paper, this species was used, to study the motion of pollen-bearing styles and the resulting pollen dispersal. *H. radicata* is a perennial plant, native to Europe, and currently is a cosmopolitan invasive species occurring in a wide range of temperate zones, including America, Japan, and Australia^17^. *H. radicata* is known to be self-incompatible. This means that the successful transport of pollen grains to different individual plants of the same species is of necessity to enable its healthy reproduction^18^. Flowering stems of *H. raticata* were collected in Kiel, Germany. They were placed in water until the youngest florets exposed fresh pollen.

### 2.2. Field observations

To observe pollen-collecting behavior of pollinators on flowerheads of *H. raticata*, we filmed videos in slow motion (120 fps) by using an iPhone 7 (Apple Inc., California, USA) together with a 30x magnifying glass (Fig. 2c-e).

### 2.3. Mechanical characterization of florets

We collected the newly opened florets from the flower heads and analyzed the morphology of the florets under a light microscope (Leica Microsystem, Wetzlar, Germany) (Fig. 1). In order to characterize the mechanical properties of the florets, we measured spring constant of the following segments: (1) styles, (2) anther tubes, and (3) filaments. Fig. 3a illustrates the experimental setup used for this purpose. Each freshly opened floret was first horizontally fixated between two wooden blocks. We fixated the floret specimens at different positions to measure the spring constant of their different segments (right-hand sketch in Fig. 3b): (1) The entire anther tube was fixated to test the style (fixation at f_1_), (2) the basal part of the anther tube including the filaments was fixated to test the anther tube (fixation at f_2_), and (3) the basal part of the petal was fixated to test the filaments (fixation at f_3_). Floret specimens were deflected using a thin metallic part mounted onto a force transducer (10g capacity; World Precision Instruments Inc., Sarasota, FL, USA). The deflections were always applied at 1 mm distal to the fixation positions at a controlled displacement speed of 0.01 mm/sec. The force required to deflect the specimens was continuously recorded using AcqKnowledge 3.7.0 software (Biopac Systems Ltd, Goleta, CA, USA). To measure the stiffness of the floret specimens, we always used the data from initial part of obtained force-displacement curves (limited to a displacement equal to 50 μm). In total, we tested 34 florets including 9 tests on styles, 10 on anther tubes, and 15 on filaments. Each floret was subjected to a single test and was not used again.

**Fig. 3.**
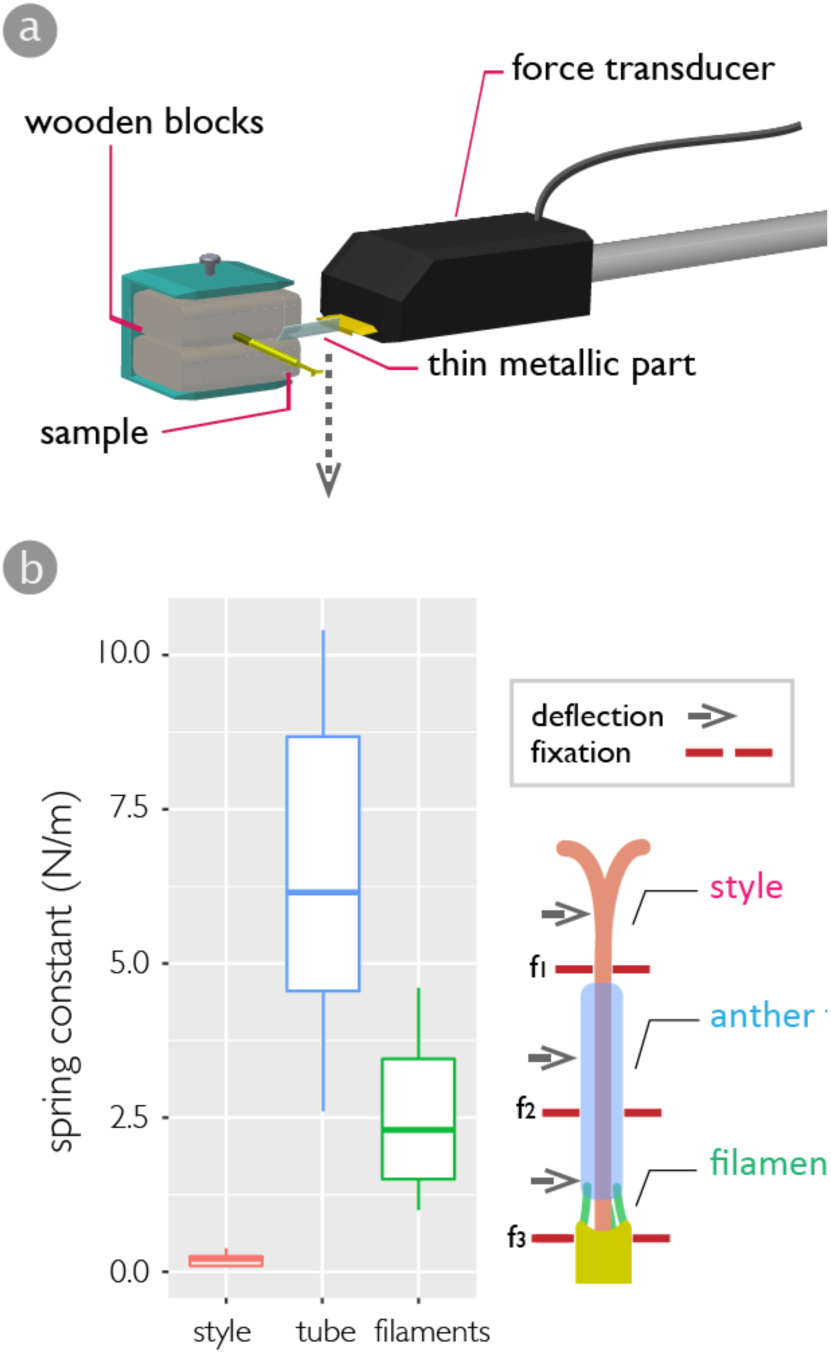
Mechanical characterization of florets. (a) The experimental setup. (b) Bar plots of the spring constant of three different segments of florets. The illustration in the right hand shows the fixations and the locations of the applied deflection. specimens were fixated at f_1_, f_2_, and f_3_ to measure the spring constants of the style, anther tube, and filaments, respectively.

### 2.4. Stylar oscillation experiments

Black circular polyethylene plates with a hole at the center were prepared for visualizing the distribution of pollen grains catapulted by stylar oscillations. Each newly opened floret was vertically fixated through the hole of the plate onto the vise to stand upright (Fig. 4). To examine whether the pollen dispersal caused by the stylar oscillation depends on the magnitude of an initial deflection, an insect pin attached to a micro-manipulator (World Precision Instruments, FL, USA) was brought in contact with one of the two following segments of the standing floret: (1) the style and (2) anther tube (Fig. 4). The contact with the anther tube resulted in large deflection of the floret, while a contact with the style caused a smaller deflection. The insect pin was kept moving horizontally until the deflected floret was released (Figs. 2i-k, 4). After the release, the floret started to oscillate and this led to the release of clumps of pollen grains, here called as ‘dispersal units’ (DU), from the style (Fig. 2j-k). The oscillations were filmed by a high-speed camera (Olympus, Tokyo, Japan) at 5000 fps and tracked using an open source tracking software (Tracker by Douglas Brown). In total, we analyzed the stylar oscillations of 55 specimens, among which 24 specimens were used to analyze the distribution and the number of pollen grains catapulted by the oscillations.

**Fig. 4.**
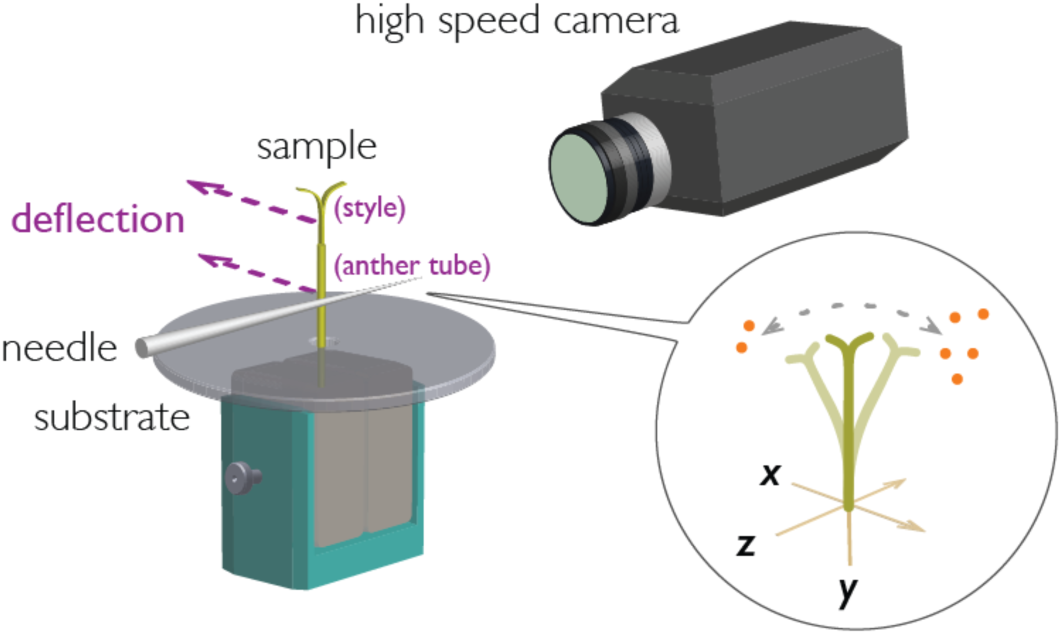
The experimental setup of the oscillation experiments. The initial deflections, which triggered the stylar oscillation, were applied to two different positions to cause two distinct deflection magnitudes. The stylar oscillations were recorded by using the high-speed camera. The dispersed pollen grains, catapulted by the stylar oscillations and landed on the circular substrates, were photographed from above.

The damping ratio, *ζ*, of a floret was obtained based on the logarithmic decrement, *δ*:

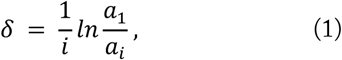

Where *a*_1_ and *a*_*i*_ are the amplitudes of the first peak and the peak, which is *i* − 1 periods away, respectively. By using eq.1, we obtained the damping ratio, *ζ*:

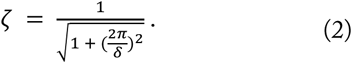

The dispersal units that landed on the plate were photographed under a microscope (Keyence, Osaka, Japan). The images were analyzed by a custom script in MATLAB (Mathworks, Natick, USA) to obtain dispersal distances. To count the number of dispersed pollen grains, the grains on the plate were collected, placed in a droplet of mineral oil on a glass slide, and sandwiched with a glass coverslip to spread out as a single layer. They were then photographed in a microscope (Leica Microsystem, Wetzlar, Germany). Using the images, the grains were counted by a custom script in MATLAB.

### 2.5. Numerical simulation of trajectories of dispersal units

Due to the technical difficulties of tracking single dispersal units based on the high-speed videos, we computed trajectories of dispersal units and their dispersal distances. During the stylar acceleration, dispersal units on the style experience inertia. They could leave the style, if the inertial force exceeds the attachment force of the dispersal units on the stylar surface. The inertial force, *F*_*i*_, required to detach a dispersal unit from the style can be obtained from:

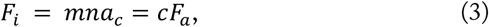

where *a*_*c*_ is the critical acceleration, *F*_*a*_ is the adhesion force of the pollen on the style, *c* is the number of contact points between a dispersal unit and the style, *m* is the mass of single pollen grains, and *n* is the number of pollen grains forming a single dispersal unit. The adhesion of pollen grains has already been measured for *H. radicata* on the stylar surface: median adhesion force, *F*_*a*_, was 98 nN (N = 50) ^16^. The mass of individual pollen grains, *m*, is 15.0 ± 0.4 ng^15^. The number of pollen grains forming a single dispersal unit, *n*, was determined by dividing the number of dispersed grains by the number of dispersal units (N = 24, *n* = 12.9 ± 5.3). The number of contact points, *c*, was set as 2 to restrict major pollen dispersal to the first cycle of oscillations, as observed in experiments. Using these data, the critical acceleration that can detach a dispersal unit from the style was calculated, and it was shown as a boxplot in Fig. 5c. In the numerical simulation, we used the median value of the critical acceleration (*a*_*c*_ = 1005 ms^−2^).

**Fig. 5.**
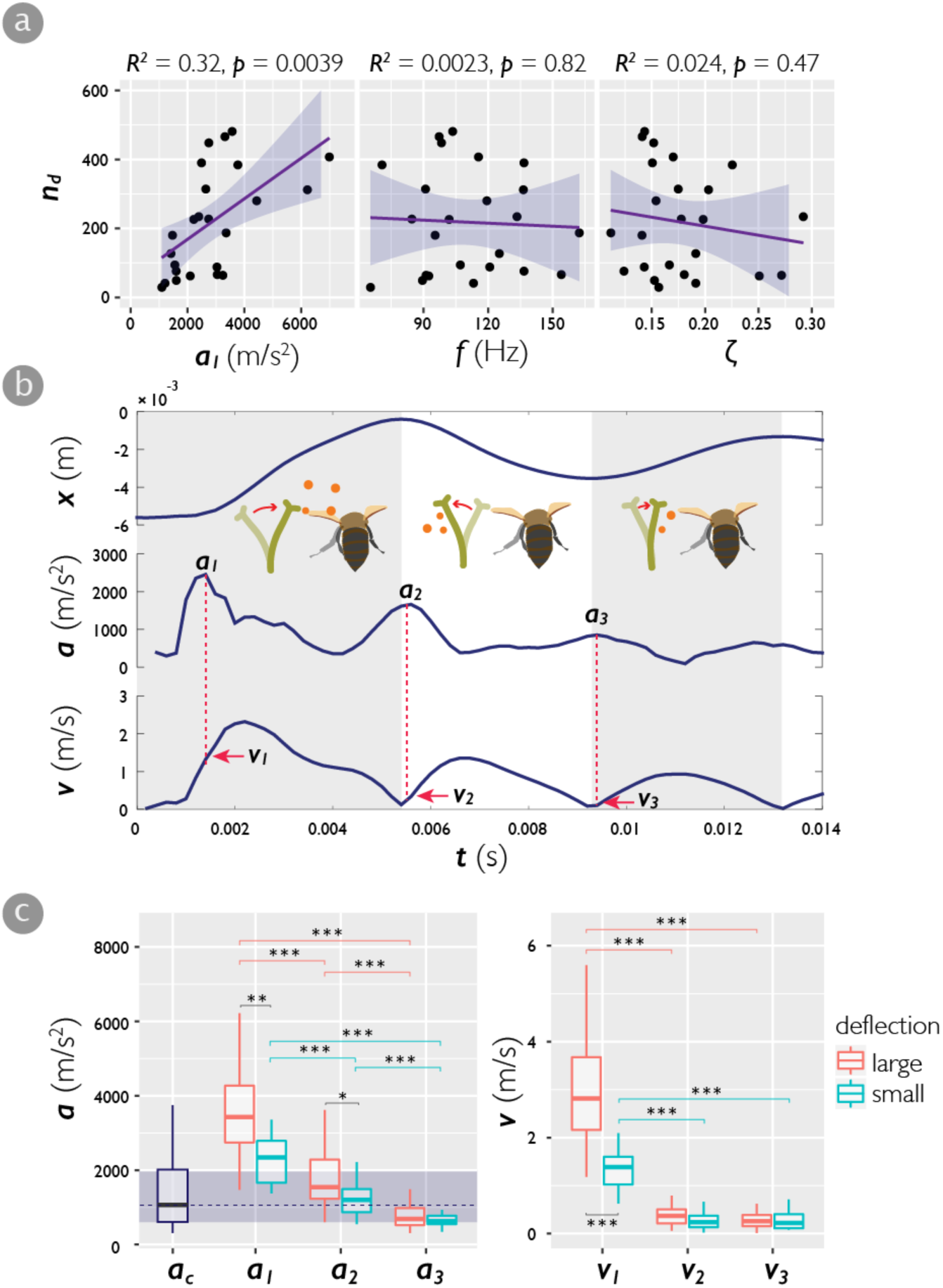
(a) Correlation analysis between the number of dispersed grains and each variable extracted from the motion data: *a*_*1*_ (left), frequency *f* (middle), and damping ratio *ζ* (right). (b) Time series of the stylar displacement *x*, linear acceleration *a*, and velocity *v*. The first, the second, and the third peak of linear acceleration were denoted as *a*_*1*_, *a*_*2*_, and *a*_*3*_, respectively. The corresponding velocities at the moment of each acceleration peak were denoted as *v*_*1*_, *v*_*2*_, and *v*_*3*_, respectively. The shadowed half cycles show the regimes where pollen should be catapulted towards a pollinator if the pollen detachment from the style occurs. (c) Bar plots of the acceleration peaks in stylar oscillations after small or large deflections (left) and the corresponding velocities at each acceleration peaks (right). The shadowed region with a vertical dashed line in the left-hand side figure show the first and the third quantile, and the median value of the critical acceleration, a*c*. The critical acceleration is required acceleration to detach dispersal units from style, calculated based on the measured pollen adhesion on the style and the average number of pollen grains in a dispersal unit (*n* = 13).

We measured the stylar velocity, *v*, and acceleration, *a*, at different time frames using the recorded high-speed videos (N = 55) (Fig. 7a, red asterisks). When the stylar acceleration was greater than the critical acceleration, the trajectory of the dispersal unit was simulated (Fig. 7a, solid curves). The initial position and velocity of each dispersal unit were set to be equal to those of the styles. The initial position of a dispersal unit (*x, y*) is referred here as the detachment point of that unit (Fig. 7b).

**Fig. 6.**
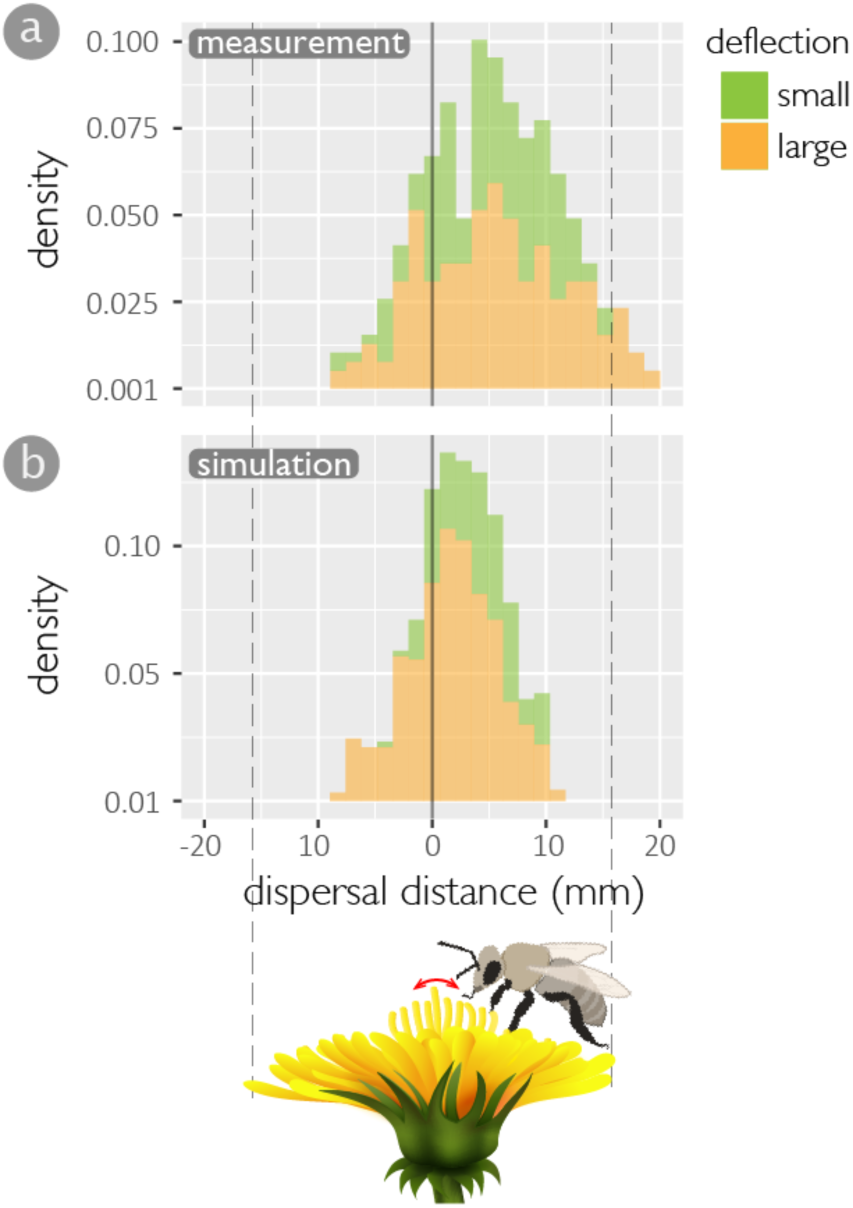
Stacked histograms of the distributions of dispersal units catapulted by stylar oscillations due to small and large deflections: (a) measurement (N = 55) and (b) simulation.

**Fig. 7.**
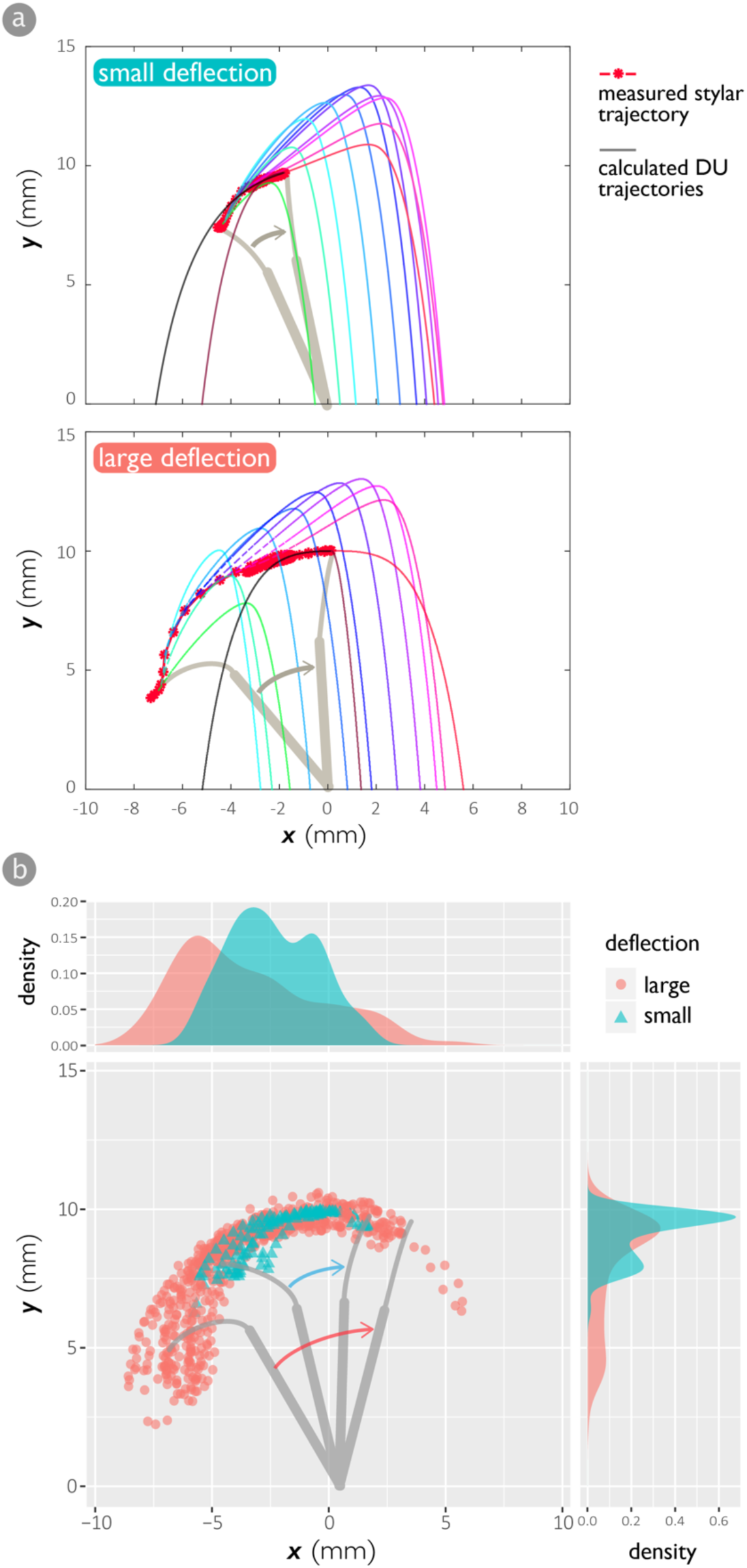
(a) Calculated trajectories of dispersal units (thin solid lines with varying colors) based on the measured trajectories of stylar oscillations (red asterisks) initiated by different deflection magnitudes. The color transition of the trajectories was intended to make them distinguishable. (b) Estimated detachment points of stylar oscillations initiated by different deflection magnitudes.

Here, we assumed the dispersal units to consist of 13 pollen grains (*n* = 13) based on the measurements, mentioned earlier. The diameter of the dispersal units can be calculated by^19^:

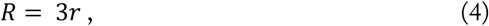

where the radius of a single pollen grain, *r*, is equal to 15 μm^15^. Reynolds number at a given time, *R*_*e*_(*t*), is given as:

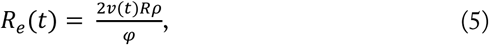

where *v*(*t*) is the velocity of a dispersal unit at a given time, *ρ* is the density of air (1.204 kg/m^3^), and *φ* is the dynamic viscosity of air at 20°C (1.825 × 10^−7^ kgm^-1^s^-1^). Considering the low range of Reynolds number (0.3 < Re < 8.5), the drag coefficient *C*_*d*_ is not constant, instead is calculated by the following empirical relationship^20^:

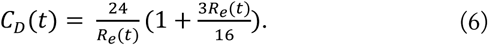

Once *C*_*d*_ is obtained, the drag force, *D*(*t*), can be calculated using the following equation in the horizontal and vertical directions, respectively:

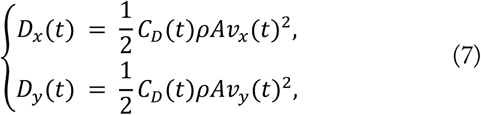

where *A* is the frontal area of the dispersal unit and equal to *πR*^2^. According to Newton’s second law, the acceleration *a*(*t*) in each direction is given as:

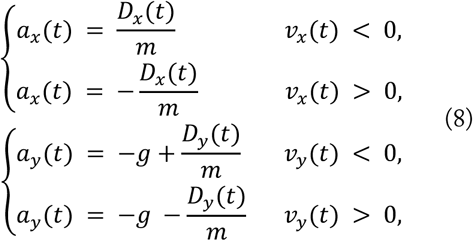

where the gravitational acceleration, *g*, is equal to 9.81 m/s^2^. The velocity and the position of the dispersal unit in the subsequent time step, *dt* = 1.0 × 10^−5^, were calculated by:

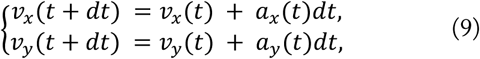

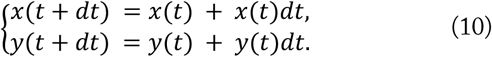

By iterating over the equations (5-10), the trajectories of dispersal units were simulated until they land on the ground, i.e. when *y*(*t*) = 0 (Fig. 7a). We assigned different colors to the trajectories of different dispersal units to make them distinguishable. The dispersal distance of a dispersal unit was defined as the landing position on the x-axis with reference to the origin, i.e. the position of the style.

We also tested the effect of the length of the style on pollen dispersal. For this purpose, we scaled the matrix of the measured stylar trajectories, *M*, using a scale factor, *s*, to obtain scaled stylar trajectories, *M*′:

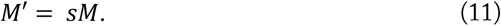

Based on the scaled stylar trajectories, *M*′, the resulting trajectories and dispersal distances of the catapulted dispersal units were simulated (Fig. 8).

**Fig. 8.**
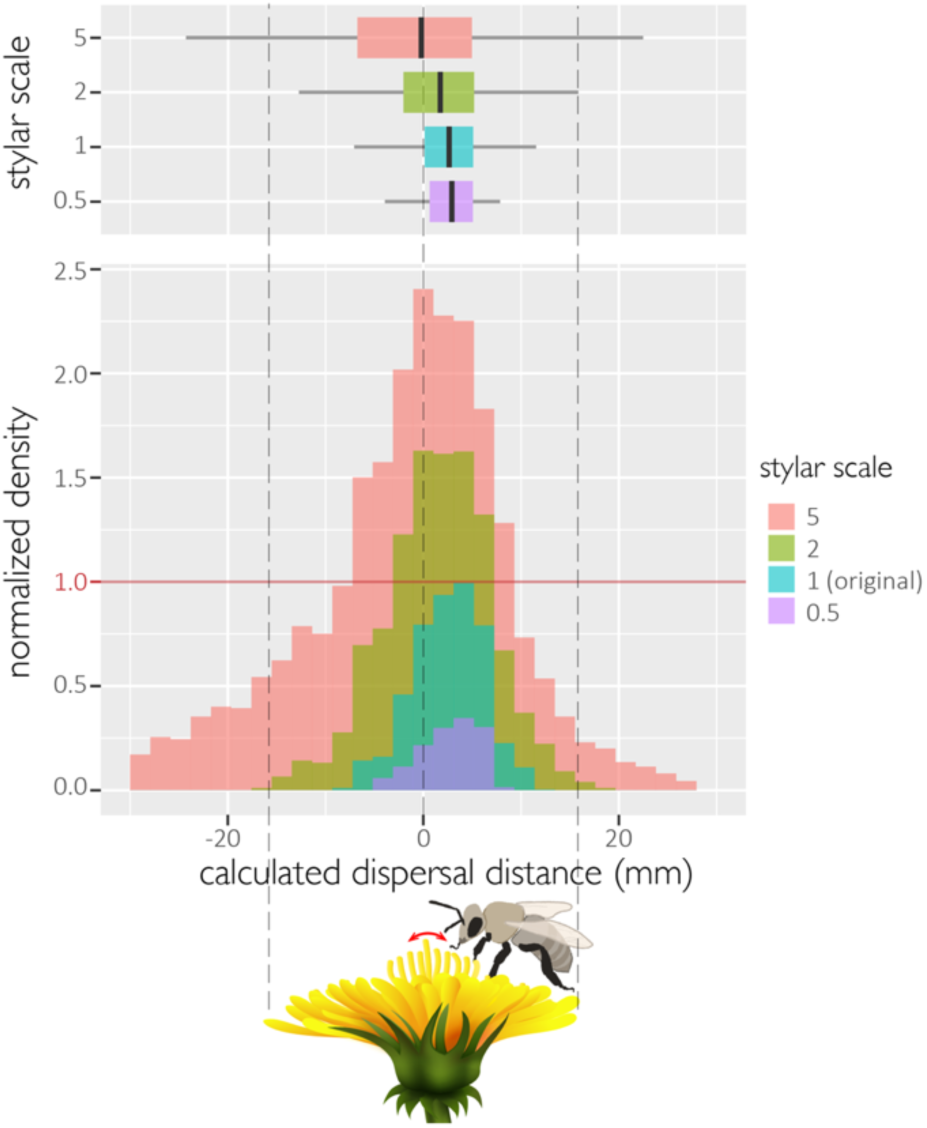
Simulated distribution of dispersal units catapulted by the oscillating styles with different lengths. The overlaid histograms are normalized to the peak density of the calculated dispersal unit distribution based on the original length of style (scale factor, *s* = 1).

## 3. Results

### 3.1. Functional segments of floret

Florets of *H. raditata* can be subdivided into three functional segments from top to bottom: an exposed distal style, an anther tube in the middle, and five basal filaments (Figs. 1, 3b). Mechanical testing of the florets revealed distinct stiffness of the three segments. The style, which passes through the anther tube and is exposed at its distal segment, has the lowest spring constant (N = 9, *K* = 0.2 ± 0.1 Nm^-1^) among other segments. It is embraced by the stiff anther tube, which features the highest spring constant (N = 10, *K* = 6.5 ± 2.7 Nm^-1^). The anther tube is connected to the petal at its proximal part by five filaments, which all together have an intermediate spring constant (N = 15, *K* = 2.3 ± 1.9 Nm^-1^). The spring constants of the three segments are significantly different from each other (Tukey multiple comparisons of means, filaments vs style: *p*-value = 0.01; anther tube vs style: *p*-value = 2.0 × 10^−7^; anther tube vs filaments: *p*-value = 1.3 × 10^−4^).

### 3.2. Mechanics of pollen dispersal

The high-speed video analysis enabled us to investigate stylar oscillations. Fig. 2i-k shows three snapshots of the stylar oscillation from release to return. When the floret was deflected, the elastic energy was mainly stored in the proximal part of the flexible style and in the filaments. However, no obvious deformation was observed in the anther tube (Fig. 2i). Upon the release, the style snapped back in the opposite direction of the applied displacement, initiating the first half cycle of oscillation. The oscillation decayed quickly, and the style returned to the resting position (N = 55, damping ratio *ζ* = 0.182 ± 0.05, frequency *f* = 119 ± 23 Hz) (Fig. 5b).

Based on the high-speed videos, we found that the first half cycle caused the major pollen dispersal (Fig. 2k). The maximum acceleration in each half cycle decreased over time (Fig. 5b and 5c, left), so that they were significantly different from each other (Tukey multiple comparisons of means, large deflections, *a*_*1*_ vs *a*_*2*_: *p*-value < 1.0 × 10^−8^, *a*_*2*_ vs *a*_*3*_: *p*-value = 0.0013, *a*_*3*_ vs *a*_*1*_: *p*-value < 1.0 × 10^−8^; small deflections, *a*_*1*_ vs *a*_*2*_: *p*-value < 2.2 × 10^−6^, *a*_*2*_ vs *a*_*3*_: *p*-value = 0.0098, *a*_*3*_ vs *a*_*1*_: *p*-value < 1.0 × 10^−8^).

As shown in Fig. 5c, for both large and small deflections, the maximum stylar acceleration in the first half cycle, *a*_1_, exceeded the critical acceleration, *a*_*c*_. However, the maximum acceleration in the subsequent half cycles (i.e. *a*_2_ and *a*_3_, respectively) largely overlapped with the critical acceleration, and the third peak, *a*_3_, mostly became lower than the median critical acceleration.

The velocity corresponding to the maximum acceleration in the first half cycle, *v*_*1*_, was by far the highest and significantly different from velocities corresponding to the accelerations in the second and third peaks, *v*_*2*_ and *v*_*3*_ (Fig. 5c, right) (Tukey multiple comparisons of means, large deflections, *v*_*1*_ vs *v*_*2*_: *p*-value < 1.0 × 10^−8^, *v*_*2*_ vs *v*_*3*_: *p*-value = 0.97, *v*_*3*_ vs *v*_*1*_: *p*-value < 1.0 × 10^−8^; small deflections, *v*_*1*_ vs *v*_*2*_: *p*-value < 2.2 × 10^−6^, *v*_*2*_ vs *v*_*3*_: *p*-value = 0.77, *v*_*3*_ vs *v*_*1*_: *p*-value < 1.0 × 10^−8^).

We examined the relationship between the number of dispersed grains, *n*_*d*_, with 3 potentially influential parameters of frequency, damping ratio, and maximum acceleration of the styles (Fig. 5a). While no correlation was found between the frequency or damping ratio with the number of dispersed grains (Pearson’s product-moment correlation, *f* vs *n*_*d*_: t = −0.22 df = 22, *p*-value = 0.82; *ζ* vs *n*_*d*_: t = −0.74, df = 22, *p*-value = 0.47), we found a significant correlation between maximum acceleration and the number of dispersed grains (Pearson’s product-moment correlation, max *a* vs *n*_*d*_: t = 3.2, df = 22, *p*-value = 0.004). The relatively low coefficient of determination (*R*^*2*^ = 0.32) can be attributed to variations in the amount of pollen grains on the examined styles.

### 3.3. Effects of deflection magnitude on pollen dispersal

Large deflections resulted in significantly higher accelerations in the first two peaks, but not in the subsequent peaks (t-test, *a*_*1*_: t = 3.90, df = 52, *p*-value = 0.00028; *a*_*2*_: t = 2.03, df = 52, *p*-value = 0.047;; *a*_*3*_: t = 0.96, df = 52, *p*-value = 0.34) (Fig. 5c, left). They also lead to significantly higher number of dispersed grains, in comparison with small deflections (t-test, t = −2.34, df = 16, *p*-value = 0.034). This is reflected by the positive correlation between the maximum acceleration in the first half cycle, *a*_*1*_, and the number of dispersed grains, *n*_*d*_, demonstrated in Fig. 5a. Whereas large deflection caused higher corresponding velocity at only the first acceleration peak (t-test, *v*_*1*_: t = 6.10, df = 52, *p*-value = 1.36 × 10^−7^; *v*_*2*_: t = 1.13, df = 52, *p*-value = 0.26; *v*_*3*_: t = 1.66, df = 52, *p*-value = 0.10) (Fig. 5c, right).

In contrast, as shown in Fig. 6a, the different deflection magnitudes had no significant effect on pollen distribution (t-test, t = −0.96, df = 397, *p*-value = 0.34). The stylar oscillation tended to catapult the dispersal units against the deflection direction with a median dispersal distance of 5.5 mm, in both large and small deflections. Surprisingly, 93.5% of the dispersal units landed within 15.8 mm from origin, which is equal to the mean radius of flowerheads of *H. radicata*.

Although our simulations underestimated the dispersal distance of dispersal units (Fig. 6), the simulated pollen distribution exhibited similar trends as the measurements: the tendency of pollen dispersal toward a pollinator and only slight effect of deflection magnitudes on the dispersal distance.

Fig. 7a shows the representative simulated trajectories of dispersal units based on the tracked stylar oscillations (red asterisks), initiated by the small and large deflections of the styles. Fig. 7b shows estimated detachment points, where the eq.3 was satisfied. Large deflections widened the range of detachment points, in comparison to small deflections. They also shifted the detachment points towards lower values in both x and y axis, compared with the small deflections (Kruskal test, x axis: chi-squared = 29.9, *p*-value = 4.65 × 10^−8^; y axis: chi-squared = 45.2, *p*-value = 1.76 × 10^−11^). It means that the larger deflections caused pollen detachment closer to the ground and further from a pollinator than the smaller deflections.

### 3.4. Effect of stylar length on pollen dispersal

We applied our numerical model to simulate pollen dispersal from styles with different lengths (*s* = 0.5, 1, 2, 5). Increasing the stylar length led to a greater number of catapulted dispersal units and a broader dispersal distance (Fig. 8). The stylar length also affected the location of the peak point in Fig. 8, meaning that the median dispersal distance shifted towards the opposite direction of a pollinator as stylar length increased (Fig. 8, bar plots).

## 4. Discussion

This study is focused on the oscillation of the style triggered by an initial deflection and its potential as a ballistic lever that catapults pollen towards a flower-visiting pollinator. Since the pollen catapult is a passive spring-driven system, the adaption to a variety of deflection magnitudes is likely to be a key to successfully deliver pollen to a pollinator and minimize waste of dispersed pollen. Here we will discuss potential role of morphological and mechanical features of a floret as well as the adhesive properties of pollen in the pollen catapult system.

### 4.1. Floret: a functional composite

Similar to other species in Asteraceae family^21^, florets of *H. raidcata* can be regarded as composites made from three morphologically distinctive segments (Figs. 1, 3b). It is known that each of the segments plays an essential role in pollen presentation^4,5,22^. Do they also contribute to the pollen catapulting towards visiting pollinators, after pollen presentation? To present pollen and successfully deliver them to visiting pollinators, the floret with the pollen-bearing style should meet these requirements: (1) high mechanical compliance during contact to store elastic energy and prevent damage and (2) catapult pollen towards a pollinator by adapting to varying magnitudes of initial deflections.

At the distal end of a floret, the flexibility of the pollen-bearing style enables high compliance under contact as well as energy storage for pollen catapult after release (Fig. 2i-k). The middle segment of the floret is reinforced by the anther tube, which functions as a support for the slender style during contacts and stabilizes the shape of the floret during non-contact periods. Previous studies have shown that the deformation of a flexible cantilever with a high aspect ratio occurs proximally to the deflection points^23^. Therefore, in the case of physical contacts at the style, the elastic energy would be stored in the broad proximal area of the floret. However, the reinforcement of the proximal segment of the floret by the anther tube prevents the deformation of this segment, limiting the amount of energy stored and released at the distal end of the style. This is likely to restrict excessive stylar acceleration, and thus to reduce the number of catapulted pollen grains.

The other energy storing segment, consisting of filaments, is situated at the base of the floret. This segment, with its intermediate stiffness, functions as a flexible joint. It enhances the mechanical compliance of the whole structure in contacts, and therefore, enables adaptation to the different deflection magnitudes. When the large initial deflection is applied, the flexible joint helps to recline the whole floret (Figs. 2i, 7a). As mentioned earlier, larger deflections lower the detachment points and shift them further from a pollinator (Fig. 7b). This is likely to counterbalance the increased initial velocity of catapulted dispersal units (Fig. 5c, *v*_*1*_) to still distribute pollen within the same range of distances, independent of the initial deflection magnitudes (Fig. 6a).

### 4.2. An optimal style length

In the plant evolution, an optimal style length has evolved. The stylar acceleration, which scales with the stylar length, dominates the pollen detachment (eq. 3, Fig. 5a). The restriction of the pollen detachment only to the first half of the oscillation cycle is a key for the directed pollen dispersal; otherwise, next oscillations can catapult pollen in the opposite direction from a pollinator, leading to an isotropic pollen distribution. This explains why increasing stylar length shifted the peak of dispersal distances against the pollinator (Fig. 8).

Furthermore, the initial velocity and height of the detached dispersal units also scaled with the stylar length, and resulted in longer dispersal distances. Considering that our simulations underestimated pollen dispersal distances, an increasing stylar length is likely to overthrow and waste more pollen grains than predicted here. On the contrary, decreased stylar length leads to the weakening of the pollen catapult with less detached pollen and shorter dispersal distances (Fig. 8, *s* = 0.5). Hence, it is likely that the length of the style is adapted to a successful delivery of pollen to a pollinator with minimizing waste of pollen.

### 4.3. Pollen adhesion

Some animals, such as insects and elks, clean themselves by vibrating their hairs to remove accumulated particles^24,25^. In such systems, natural frequency governs particle detachment:

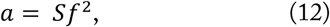

where *a* is the acceleration, *S* is the spacing between the hairs, which dictates the maximum deflection of the hairs, and *f* is the natural frequency^24^. However, here, we did not find correlations between the natural frequency and the number of dispersed grains (Fig. 5a) nor between the natural frequency and maximum linear acceleration (Fig. S1).

It is important to mention that the major pollen detachment occurs only at the first half oscillation cycle. Hence, the general oscillatory frequency has very little influence on the pollen detachment. To restrict the pollen detachment only to the first half cycle, the critical acceleration, *a*_*c*_, should be less than the first maximum acceleration, *a*_1_, but greater than maximum accelerations in any other oscillation cycles (Fig. 5c, left).

As shown in eq.3, the critical acceleration depends not only on the pollen-style adhesion, but also the pollen-pollen adhesion, which governs the number of pollen grains in a dispersal unit. The weaker pollen-pollen adhesion, therefore, decreases the number of grains in a dispersal unit, in comparison to *n* = 13 measured here. Since the inertial force is proportional to the mass of the dispersal unit, a smaller number of pollen in a dispersal unit can challenge its detachment from the style. If the number of grains is as small as *n* = 3, the median critical acceleration would be greater than most of the measured maximum stylar accelerations, and this would completely emasculate the pollen catapult. In contrast, higher pollen-pollen adhesion would result in higher number of grains in the dispersal units. This would, on the other hand, lead to a prolonged, and probably, an isotropic pollen dispersal (both towards and against the pollinator) and, therefore, increase the number of wasted pollen. Hence, for a guided pollen dispersal towards a pollinator, pollen adhesion and the size of dispersal units should be kept in a certain range.

Unlike wind-pollinated pollen, the insect-pollinated ones are covered by a thick layer of a viscous oily substance called pollenkitt^26^, which helps to form large dispersal units by bonding pollen grains together^26,27^. In addition to its general function as “pollen adhesive”, we recently showed that pollenkitt inhibits pollen adhesion by weakening water capillary attraction^15^. Therefore, we can assume that pollenkitt may play a role in maintaining pollen adhesion in a specific range. Future studies are needed to quantitatively investigate the role of pollenkitt in pollen-pollen adhesion.

### 4.4. Phase shift in the first half oscillation cycle

Once the pollen detachment occurs, the pollen dispersal is mainly governed by the initial velocity of the dispersal units, rather than their acceleration. Hence, the velocity corresponding to each maximum stylar acceleration (Fig. 5c, right) is another key for the guided pollen dispersal towards a pollinator.

Unlike the stylar acceleration peaks, the corresponding stylar velocities abruptly decayed, so that the velocities corresponding to the second and third maximum accelerations (i.e., *v*_*2*_ and *v*_*3*_, respectively) were not significantly different from each other (Fig. 5c, right). Only the first corresponding velocity, *v*_*1*_, is significantly higher than the others, because of the phase shift between the stylar acceleration and velocity at the first half cycle.

In a free oscillation, the displacement, *x*(*t*), is given as:

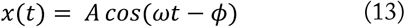

where *A* is an arbitrary constant and *ω* is the angular frequency. Differentiating eq.13 with respect to *t* gives the velocity *v*(*t*) as:

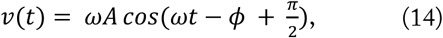

In the same manner, the acceleration, *a*(*t*), is given as:

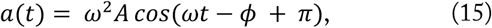

The phase difference between the acceleration and velocity is equal to 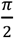, or a quarter of an oscillation cycle. Therefore, in the free oscillation, the corresponding velocities at the moment of acceleration peaks, ideally, are the local minima. As shown in Fig. 5b, this typical phase difference appeared in the stylar oscillations, resulting in the extremely small values of *v*_*2*_ and *v*_*3*_. However, the first corresponding velocity, *v*_*1*_, was shifted from the local minimum; therefore, *v*_*1*_ was by far the highest than other corresponding velocities. The cause of the phase shift is seemingly due to the style sliding over the insect pin during the release, and therefore, a detailed study on this topic will be required in the future.

### 4.5. Biological significance

Asteraceae are known for attracting various pollinators^6^. Their robust adaptability makes a number of them notorious invasive species^28,29^. To enable their versatile reproduction, they are likely to use multiple strategies to secure pollen transfer to diverse pollinators. Each pollinator species may have its own difficulties to be a functional pollen vector for the benefit of plants. Bees, for example, are both pollen-transporting vectors and pollen robbers. Actively collected pollen grains, are soaked with their saliva and are packed in a specialized pollen-carrying apparatus, and no longer available for pollination^30,31^. Some pollinators, such as flies from the family Bombyliidae, with their specialized long mouth parts and slender limbs less frequently make physical contacts with styles on flowers, yet are able to steal nectary rewards^32^ (Fig. 2f-h).

In this paper, we discussed that the superficially unremarkable floret of *H. radicata*, by optimizing its (1) morphological and mechanical properties, (2) length, (3) adhesive properties of pollen, and (4) phase difference between the stylar acceleration and velocity, functions as a projectile tool to catapult pollen towards pollinators. The airborne pollen dispersal system is advantageous to randomly deliver pollen to blind spots of pollinators as well as to engage the long-limbed insects beyond the physical reach of the styles^13,14^. It is also remarkable that more than 90% of the dispersal units landed within a region with the same size as that of the flowerhead. Therefore, even if the dispersal units fail to reach the target pollinator, they still remain within the flowerhead, waiting for being transferred to the forthcoming pollinators.

It is well known that florets of Asteraceae consecutively mature from the outer row towards the central row. For many species of them, including *H. radicata*, this means that young and most pollen-bearing styles are situated in the center of the flowerhead^33^. The tuned travel distance of the dispersal units, therefore, may explain the order of the floret maturity in Asteraceae.

Since the working principles of the pollen catapult arises from the common floral feature of Asteraceae, the airborne pollen delivery system potentially contributes to the versatile adaptivity of the most successful flowering plant family.

## 5. Conclusions

In this study, we suggested novel pollen dispersal strategy resulted from the stylar oscillation of *H. radicata*. Based on the combination of high-speed motion analysis, mechanical tests, and numerical simulations, we found that the morphologically and mechanically distinctive segments composing the floret, as well as the optimal pollen adhesion, contribute to (1) catapulting pollen towards a visiting pollinator while (2) minimizing the wasteful pollen dispersal by (3) adapting to different amount of oscillation-inducing external deflections. Since the ballistic pollen dispersal, arises from a standard floral morphology of Asteraceae, it is likely to play a role in the reproductive assurance of the most ubiquitous flowering family on the planet.

## 6. Acknowledgements

The authors thank Alexander Köhnsen for the help in motion tracking, and the Youtube channel “JustNature” for the permission to use a video-footage of a Bombyliidae. Yuka Ito is acknowledged for the permission to use her iPhone for a field work, and Aria Ito for an inspiration of this study.

## 8. Author contributions

S.I., H.R., and S.N.G. conceptualized the study. S.I. and H.R. designed the experiments. S.I. conducted the research, collected and analyzed the data. S.I. and H.R. participated in the data presentation. S.N.G. provided equipment and supervised the study. S.I. wrote the first draft. H.R. and S.N.G. edited the manuscript. All authors discussed the results and gave the final approval for publication.

