## Supplementary material for "A ballistic pollen dispersal strategy hidden in stylar oscillation": Fig. S1

Supplementary Figures

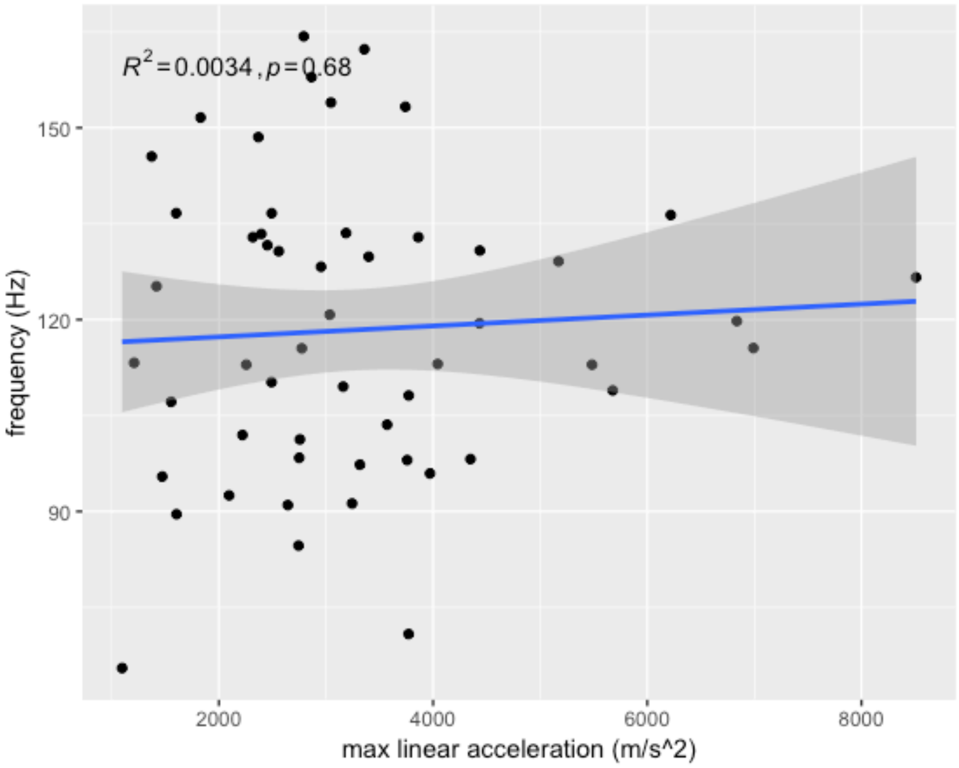


Fig. S1 Correlation analysis between frequency *f* and maximum linear accelerations (max *a*).
